# The 3D construction of leaves is coordinated with water use efficiency in conifers

**DOI:** 10.1101/2021.04.23.441113

**Authors:** Santiago Trueba, Guillaume Théroux-Rancourt, J Mason Earles, Thomas N. Buckley, David Love, Daniel M Johnson, Craig Brodersen

**Author notes:** Author for correspondence: Santiago Trueba.

## Abstract

- Conifers prevail in the canopies of many terrestrial biomes, holding a great ecological and economic importance globally. Current increases in temperature and aridity are imposing high transpirational demands and resulting in conifer mortality. Therefore, identifying leaf structural determinants of water use efficiency is essential in predicting physiological impacts due to environmental variation.
- Using synchrotron-generated microCT imaging, we extracted leaf volumetric anatomy and stomatal traits in 34 species across conifers with a special focus on *Pinus*, the richest conifer genus.
- We show that intrinsic water use efficiency (WUE_i_) is positively driven by leaf vein volume. Needle-like leaves of *Pinus*, as opposed to flat leaves or flattened needles of other genera, showed lower mesophyll porosity, decreasing the relative mesophyll volume. This led to increased ratios of stomatal pore number per mesophyll or intercellular airspace volume, which emerged as powerful explanatory variables, predicting both stomatal conductance and WUE_i_.
- Our results clarify how the three-dimensional organization of tissues within the leaf has a direct impact on plant water use and carbon uptake. By identifying a suit of structural traits that influence important physiological functions, our findings can help to understand how conifers may respond to the pressures exerted by climate change.

## Introduction

Conifer forests thrived on the Earth surface during the Mesozoic until the radiation and diversification of angiosperms during the Cretaceous, which was followed by angiosperm ecological dominance attributed to increased physiological performance and reduced generation times (Bond, 1989; Boyce *et al*., 2009; Crepet & Niklas, 2009; de Boer *et al*., 2012). Yet, after 100 million years of competition with the angiosperms, conifers remain prominent in the canopy of many biomes (Brodribb *et al*., 2012). Conifers are found in ecosystems from high latitudes to the equator, and they have a major economic importance in the wood and paper industries (McFarlane & Sands, 2013). Conifer-dominated forests are not exempt from the impacts of drought and aridity resulting from ongoing global climatic changes (Dai, 2011; Brodribb *et al*., 2020; Kharuk *et al*., 2021). This is particularly alarming given that 48% of the 722 conifer taxa of the world are currently threatened (Blackmore *et al*., 2011). Global increases in temperature coupled with rising vapor pressure deficit (VPD) place increased strain on plant hydraulic and photosynthetic systems (Choat *et al*., 2018; Grossiord *et al*., 2020). There is strong evidence that tree water use efficiency (WUE) has increased in recent decades, most likely the result of rising atmospheric CO_2_, allowing plants to open their stomata less frequently, thereby conserving water (Keenan *et al*., 2013; Mathias & Thomas, 2021). However, the underlying physiological mechanisms behind this trend need to be further elucidated (Guerrieri *et al*., 2019). Furthermore, there is still a lack of knowledge about how leaf structural organization influences key functions such as photosynthetic carbon acquisition, stomatal conductance, and the interplay between both driving intrinsic water use efficiency (WUE_i_).

Leaf-level photosynthetic metabolism has an important role in maintaining global ecological processes (Hetherington & Woodward, 2003). Therefore, exploring tissue organization inside the leaf, and specifically the mesophyll cells where water and gas diffusion occurs, is important for understanding carbon, water, and energy fluxes at whole ecosystem levels. The leaf mesophyll consists of photosynthetic parenchyma cells located between the epidermis and the bundle sheath layers surrounding the veins. Once inside the leaf, the diffusion of CO_2_ through the intercellular airspace (IAS) and to the chloroplasts (i.e. mesophyll conductance) is a major constraint on photosynthetic performance (Flexas *et al*., 2012; Gago *et al*., 2020). This pathway includes the IAS, but also the diffusion of CO_2_ across mesophyll cell walls, cell membranes, and the chloroplast envelope, which can significantly limit photosynthesis in gymnosperms (Carriquí *et al*., 2020). Previous work has shown that mesophyll surface area per leaf area (S_m_ ; µm^2^ µm^-2^) had a strong influence on maximum photosynthesis (Nobel *et al*., 1975; Smith & Nobel, 1977). More recently, it has been suggested that the surface area of the mesophyll exposed to the IAS per unit of mesophyll volume (SA_*mes*_ /V_*mes*_ ; µm^2^ µm^-3^) can influence plant photosynthetic performance given that the mesophyll-IAS boundary is the primary interface between the atmosphere and the photosynthetic cells (Earles *et al*., 2018; Théroux-Rancourt *et al*., 2021). Other volumetric anatomical traits such as mesophyll porosity (i.e. IAS as a fraction of total mesophyll volume; µm^3^ µm^-3^) might be relevant in promoting diffusion through the IAS, and it has been suggested that mesophyll palisade porosity is correlated to stomatal conductance across four different *Arabidopsis* mutants (Lundgren *et al*., 2019). The relevance of such anatomical traits arises from the hypothesis that photosynthetic capacity could be enhanced by increasing surface area of mesophyll exposed to the IAS, allowing for more potential surface for CO_2_ diffusion across mesophyll cell walls, and chloroplast envelopes (Evans *et al*., 2009; Earles *et al*., 2018). However, the correct estimation of such anatomical traits using standard two-dimensional (2D) techniques is difficult since it relies on 2D approximations of the complex, three-dimensional (3D) shape of the mesophyll (Théroux-Rancourt *et al*., 2017; Earles *et al*., 2019).

Given the elementary vascular architecture of the conifer leaf, which typically consists of a single vein without further branching (Zwieniecki *et al*., 2004; Brodribb *et al*., 2010), and the large variation in conifer leaf shape (Fig. 1), area-based traits might not allow for an accurate comparison of strategies to optimize leaf structure with function within conifers (Earles *et al*., 2019; Théroux-Rancourt *et al*., 2021). Further, given the structural relevance of the central vasculature, traits other than those related to the mesophyll might also influence CO_2_ and water diffusion in the conifer leaf. Therefore, a central question is whether non-vascular tissues are tightly coupled to vein volume with proportional relationships, or whether they show greater variability to compensate for reduced hydraulic efficiency. We predict that higher volumes of vascular tissue per unit leaf volume (V_*vein*_ /V_*leaf*_ ; µm^3^ µm^-3^) should be a good predictor of gas exchange and WUE_i_ because of the mechanistic link between hydraulic conductance and maximum photosynthetic rates in conifers (Brodribb & Feild, 2000). We also predict a positive relationship between gas exchange efficiency and the ratio of mesophyll surface area exposed to the IAS and vein volume (SA_*mes*_ /V_*vein*_ ; µm^2^ µm^-3^), where a greater investment in vein volume per unit area of bulk tissue surface should increase the hydraulic capacity to replace water lost to transpiration. In conifer leaves, water moves across the bundle sheath and the transfusion tissue before reaching the mesophyll (Hu & Yao, 1981). Therefore, bundle sheath and transfusion tissue volume relative to total leaf volume (V_*BS*+*TT*_ /V_*leaf*_ ; µm^3^ µm^-3^), should also influence the efficiency of water movement within the leaf. Along with the previously described features, many conifer species also possess resin ducts, which play a major role in chemical and physical defense (Gaylord *et al*., 2013; Breshears *et al*., 2018), but necessarily displace vascular or photosynthetic tissue, incurring both maintenance and construction costs, but also lost opportunity costs for net carbon gain. Finally, water movement inside the leaf ends at the stomatal pores, which play a major role in regulating water loss and maintaining water status in conifers (Brodribb *et al*., 2014). Within this context, it is possible to determine the proportionality of different anatomical traits, the coordination between supply and demand for CO_2_ and H_2_O, and the physical constraints of leaf construction. To probe these relationships, we describe how conifers build their elementary, yet diverse leaves, and how structural features relate to key physiological traits.

**Figure 1.**
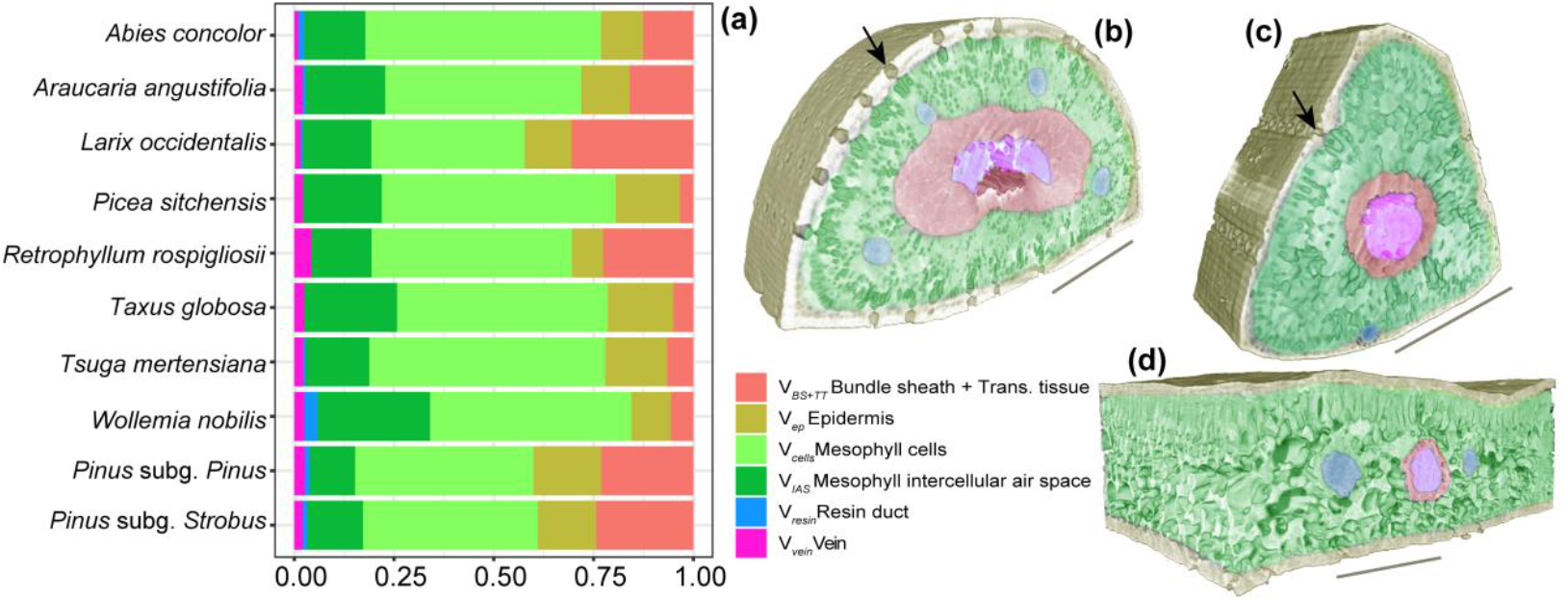
Proportion of different tissue volumes relative to total leaf volume inside the 3D leaf space in conifers (a). Average values of *Pinus* species belonging to the *Pinus* subgenus (21 spp) and *Strobus* (5 spp) are included. MicroCT 3D images of needle-leaved *Pinus pungens* (b; *Pinus* subgenus) and *Pinus monticola* (c; *Strobus* subgenus) and flat-leaved *Wollemia nobilis* (d). Segmented tissues are indicated with different colors and stomatal apertures are indicated with arrows. Scale bars = 250 µm. A plot of the relative tissue volume of all 34 species is available in Fig. S2.

This study presents a survey of the three-dimensional organization of the conifer leaf using microCT imaging (Table S1). Our study includes 34 conifer species (Fig. S1), with a special focus on the genus *Pinus*, the largest extant genus of conifers (Gernandt *et al*., 2005). *Pinus* species can be found in a broad range of environmental conditions, suggesting wide structural and functional diversity (Martínez-Vilalta *et al*., 2004). *Pinus* subgenera *Pinus* and *Strobus* can be distinguished by having two or one vascular bundles respectively, centrally located inside the needle-like leaf (Gernandt *et al*., 2005) (Fig. 1a-c). Despite their rather simple anatomical organization, conifer leaves have extensive morphological diversity ranging from flat leaves to needle-like leaves with different degrees of transversal flattening. Leaf morphological diversity in conifers results in different physiological performances, with flat-leaved species having lower photosynthetic assimilation and respiration rates than needle-leaved species (Brodribb & Feild, 2008; Schmiege *et al*., 2020; Schmiege *et al*., 2021). Yet, the differences in 3D anatomical structure across conifers with different leaf morphologies, which could explain their contrasting photosynthetic performance, need to be elucidated. We hypothesize that features enhancing mesophyll surface area for gas diffusion, will be positively correlated with the light-saturated assimilation rate of CO_2_ (*A*_sat_) and maximal stomatal conductance (*g*_smax_). We also provide a volume-based stomatal density estimation, a trait we expect better captures the interplay between the evaporative surfaces invested in the non-laminar mesophyll volume and the number of stomata needed to provide CO_2_, with the expectation that leaves with higher number of stomata per mesophyll volume would have both higher rates of gas exchange and WUE_i_ due to an enhancement of the epidermal pores serving as evaporative surface relative to the photosynthetic tissue.

## Materials and Methods

### Plant material

Sampling included 34 conifer species from various biomes, physiologies, and leaf morphologies. Sampling included taxa from four different families of conifers: Araucariaceae (2 spp), Pinaceae (30 spp), Podocarpaceae (1sp), and Taxaceae (1 sp). Our sampling particularly focused on the genus *Pinus* with 26 species including representatives from the two subgenera: *Pinus* (21 spp) and *Strobus* (5 spp), which differ in the number of vascular bundles per leaf (Gernandt *et al*., 2005). Sampling also represents three distinct conifer leaf morphologies: flat leaves (*Araucaria, Retrophyllum, Taxus* and *Wollemia*; Fig. 1d), flattened needles (*Abies, Larix, Picea* and *Tsuga*), and needle-like leaves (*Pinus*; Fig. 1b,c). Flattened needles, such as those commonly found in non-*Pinus* Pinaceae, are generally shorter and flattened in cross-section as compared to *Pinus* needle-like leaves, which have almost equal width and height (Fig. 1b,c) and are generally longer. Fully expanded leaves from adult plants were collected in the Berkeley Arboretum of the University of California Botanical Garden, and the University of Georgia’s Thompson Mills Arboretum. Samples from both locations were used for microCT scanning and gas exchange measurements. Whole shoots were cut, wrapped in moist paper towels, and transported in dark plastic bags to avoid desiccation before scanning.

### X-ray microtomography (microCT) scanning and image segmentation

MicroCT imaging was performed at the Lawrence Berkeley National Laboratory Advanced Light Source, beamline 8.3.2. Leaves were scanned within 24h of excision. Samples were wrapped with a polyimide (Kapton) tape, which allows x-ray transmittance while preventing sample desiccation. Wrapped leaf samples were placed in a plastic 1000 µL pipette tip with the lower end submerged in water and centered in the microCT x-ray beam. Scans were completed in c. 15 minutes in continuous tomography mode at 21 keV capturing 1,025 projection images of XYZ ms each. Images were captured using alternatively 5x or 10x objective lenses depending on leaf diameter, yielding final pixel resolutions of 1.27 μm and 0.625 μm. Images were reconstructed using TomoPy (Gürsoy *et al*., 2014). Raw tomographic datasets were reconstructed using both gridrec and phase retrieval methods, both of which yield complementary results being efficient in segmenting cell boundaries and larger air voids, respectively (Théroux-Rancourt *et al*., 2017). Image stacks of c. 2600 8-bit grayscale images were generated from the reconstruction process. Airspace was segmented in both gridrec and phase reconstruction methods by visually defining a range of pixel intensity values and the binary image stacks from both reconstruction methods were combined. Boundaries delimiting the areas occupied by the bundle sheath + transfusion tissue, epidermis, mesophyll, resin ducts, and veins were manually drawn using a graphic tablet (Wacom Cintiq Pro 16, Wacom Co, Saitama, Japan) in ImageJ (Schneider *et al*., 2012). Leaf veins were depicted here as the vascular bundle comprising both xylem and phloem tissues. Leaf tissue boundaries were drawn as regions of interest (ROIs) in six to eight images randomly distributed across the full stack. The combination of the binary image derived from both reconstruction methods, along with the tissue boundaries, resulted in a composite image stack where each leaf tissue was classified. Leaf segmentation, which allowed us to automatically delimit different tissues across the full stack using a limited set of hand-segmented composite slices was done using random-forest classification (Théroux-Rancourt *et al*., 2020).

### Three-dimensional anatomy and stomatal traits

We extracted the volume and surface area of leaf anatomical traits from the full segmented stacks (Théroux-Rancourt *et al*., 2020). We estimated the volumes of the epidermis (V_*ep*_), bundle sheath and transfusion tissue (V_*BS*+*TT*_), mesophyll cells (V_*cell*_), mesophyll intercellular airspace (V_*IAS*_), resin ducts (V_*resin*_), and veins (V_*vein*_). Mesophyll volume (V_*mes*_) was estimated as the sum of V_*cell*_ and V_*IAS*_. All volume metrics are reported in μm^3^. Relative volumes for each tissue (in μm^3^ μm^−3^) were estimated as a fraction of tissue per V_*leaf*_, total leaf volume. We calculated mesophyll porosity (μm^3^ μm^−3^) as V_*IAS*_ /V_*mes*_. The mesophyll surface area exposed to the intercellular airspace (SA_*mes*_) was used to estimate the mesophyll surface area per mesophyll volume (SA_*mes*_ /V_*mes*_ ; μm^2^ μm^−3^). Additionally, we estimated the exposed mesophyll surface area per bundle sheath volume (SA_*mes*_ /V_*BS*+*TT*_ ; μm^2^ μm^−3^) and vein volume (SA_*mes*_ /V_*vein*_ ; μm^2^ μm^−3^). Total leaf area A_*leaf*_ (μm^2^) was measured by summing up the perimeter of each slice and multiplying it by slice depth. We used the ratio SA_*mes*_ /A_*leaf*_ to calculate the mesophyll surface area per total leaf area (S_m_ ; μm^2^ μm^−2^). Stomatal estimations were performed by counting all visible stomata on the leaf surface of each scan using Avizo 9.4 software (FEI Co. Hillsboro, OR, USA). Absolute stomatal counts were used in relation to mesophyll volumetric anatomy to estimate traits accounting for the interaction of stomata pore number and V_*IAS*_, V_*mes*_, and SA_*mes*_ units. Mesophyll volumetric features were assessed in mm^3^ for stomatal pore density estimations. We also accounted for stomatal number per leaf surface to assess potential differences in stomatal density estimations based on surface vs volume fractions. Further, we performed a comparison of surface- and volume-based anatomical estimations using 2D slices and the full 3D stack. Methods are further explained in Notes S1. Average trait values for each species are available in Dataset S1. A list of measured anatomical variables, including abbreviations and units, is available in Table S1.

### Gas exchange and water use efficiency measurements

Maximum stomatal conductance (*g*_smax_ ; mmol m^− 2^ s^− 1^) and light-saturated CO_2_ assimilation rate (*A*_sat_ ; µmol m^− 2^ s^− 1^) were measured on a subset of 18 species (Dataset S1) and used to estimate leaf-level intrinsic water use efficiency (WUE_i_ = *A*_sat_ /*g*_smax_ ; µmol mol^− 1^). *A*_sat_ data for *Pinus strobus* was removed prior to analyses due to potential measurement inaccuracies. Gas exchange measurements were performed with a LICOR-6800 gas exchange system (LI-COR biosciences, Lincoln, NE, USA) between 9:00 and 14:00 on fully sunlit outer canopy foliage. Chamber temperature was set to 25° C, light source was set to 1500 µmol m^− 2^ s^− 1^ and chamber CO_2_ was set to 400 ppm. Following the gas exchange measurements, the leaf area contained within the chamber was marked using a permanent marker, collected, and the projected leaf area was measured using a leaf area meter (LI3100C, LI-COR biosciences, Lincoln, NE, USA).

### Data analysis

All statistical analyses and data treatment were performed using R v.3.6.3 (R Core Team, 2020). Anatomical traits were assembled and averaged for each species for data analysis. Assumptions for residual homogeneity and normality were tested prior to data analyses. Phylogenetic relationships, including branch length calibrations and divergence times, were obtained from published data (Magallón *et al*., 2015; Smith & Brown, 2018). To predict leaf physiological traits based on volumetric anatomical variables we used phylogenetic generalized least-squares analyses (PGLS) with a lambda (λ) maximum likelihood optimization to control for phylogenetic non-independence between related species (Felsenstein, 1985; Freckleton *et al*., 2002). A two-parameter exponential function, *y* = *a*(1−exp(−*bx*)), was additionally employed to describe the relationship of WUE_i_ and V_*vein*_ /V_*leaf*_ (*a* = 130, *b* = 109, *r*^2^=0.50). The assemblage of the composite phylogenetic tree of studied conifers (Dataset S2) was carried out using the package ‘ape’ (Paradis & Schliep, 2018) and PGLS models were fit using the package ‘caper’ (Orme *et al*., 2018). Standard major axis (SMA) were implemented with the package ‘smatr’ (Warton *et al*., 2012) to test allometric scaling between tissue volumes. A principal component analysis (PCA) was used to explore the covariation of selected traits and the distribution of leaf morphologies and conifer groups as explained by the physiological and anatomical traits measured. Given the marked differences in leaf anatomical structure across conifer leaf morphological types, and between *Pinus* subgenera *Pinus* and *Strobus*, we explored potential variation in 3D anatomical traits by plotting each group within the PCA. We further explored differences across leaf morphologies and between conifer clades by performing a one-way ANOVA on measured structural traits. Physiological features were not compared between leaf morphological types due to insufficient data availability. A similar variance meta-analysis, including post hoc Tukey’s honest significant differences, was employed to compare the studied conifer species with other gymnosperms, along with angiosperms and ferns. Data for comparisons across plant groups was obtained from a recently published study (Théroux-Rancourt *et al*., 2021) and it is available in Dataset S3.

## Results

The mesophyll (V_*mes*_ /V_*leaf*_), including cells and airspace, represents the dominant leaf volume fraction for all 34 measured conifer species in this study, occupying an average of 60% of the total leaf volume (Fig. 1; Fig. S2). The second largest volume fraction was either the combined bundle sheath and transfusion tissue (V_*BS*+*TT*_ /V_*leaf*_) that surrounds the vascular cylinder, or the epidermis (V_*ep*_ /V_*leaf*_), which represented an average of 22% and 16% of the leaf volume, respectively (Fig. 1a). Veins (V_*vein*_ /V_*leaf*_) and resin ducts (V_*resin*_ /V_*leaf*_) represented the smallest fraction of the total leaf volume (Fig. 1a), and resin ducts were completely absent in six measured species (Dataset S1). We found weak structural coordination amongst tissue volumes within the conifer leaf (standard major axes; Table S2). However, a negative allometric scaling between V_*ep*_ /V_*leaf*_ and V_*mes*_ /V_*leaf*_ was observed (*r*^2^ = 0.27, *p* < 0.01), indicating that increasing the relative allocation to the mesophyll was done in conjunction with a decrease in the relative allocation to the epidermis. Average values of all measured volumetric traits for each species are included in Dataset S1.

A multivariate analysis of trait covariation defined two major axes explaining 48% of inertia (Fig. 2). Inertia of the first axis was mainly explained by V_*IAS*_ /stomate, the amount of air volume connected to a stomate (18.27%), stomata/V_*mes*_, the stomatal density per mesophyll volume (15.04%), V_*mes*_ /V_*leaf*_ (14.74%) and *g*_smax_ (14.08%). Increasing *g*_smax_ was associated on the first axis with increases in stomata/V_*mes*_ and *A*_sat_, and decreases in mesophyll porosity, V_*mes*_ /V_*leaf*_, and V_*IAS*_ /stomate (Fig. 2). The second axis was largely explained by V_*vein*_ /V_*leaf*_ (18.04%), SA_*mes*_ /V_*vein*_ (14.83%) and S_m_ (14.45%). Increasing WUE_i_ was associated on the second axis with increases in S_m_ and V_*vein*_ /V_*leaf*_, and decreases in stomata/V_*mes*_ (Fig. 2). Leaf morphological types were dispersed along both major axes (Fig. 2). Needle-like leaves were isolated due to their higher V_*ep*_ /V_*leaf*_ and stomata/V_*mes*_. Flat leaves and flattened needles largely overlapped due to their porous and voluminous mesophylls as opposed to needle-like leaves (Fig. 3a,b). Flat leaves and flattened needles also converged in having lower stomata/V_*mes*_ (Fig. 3c) and higher V_*IAS*_ /stomate (Fig. 3d) than needle-like leaves.

**Figure 2.**
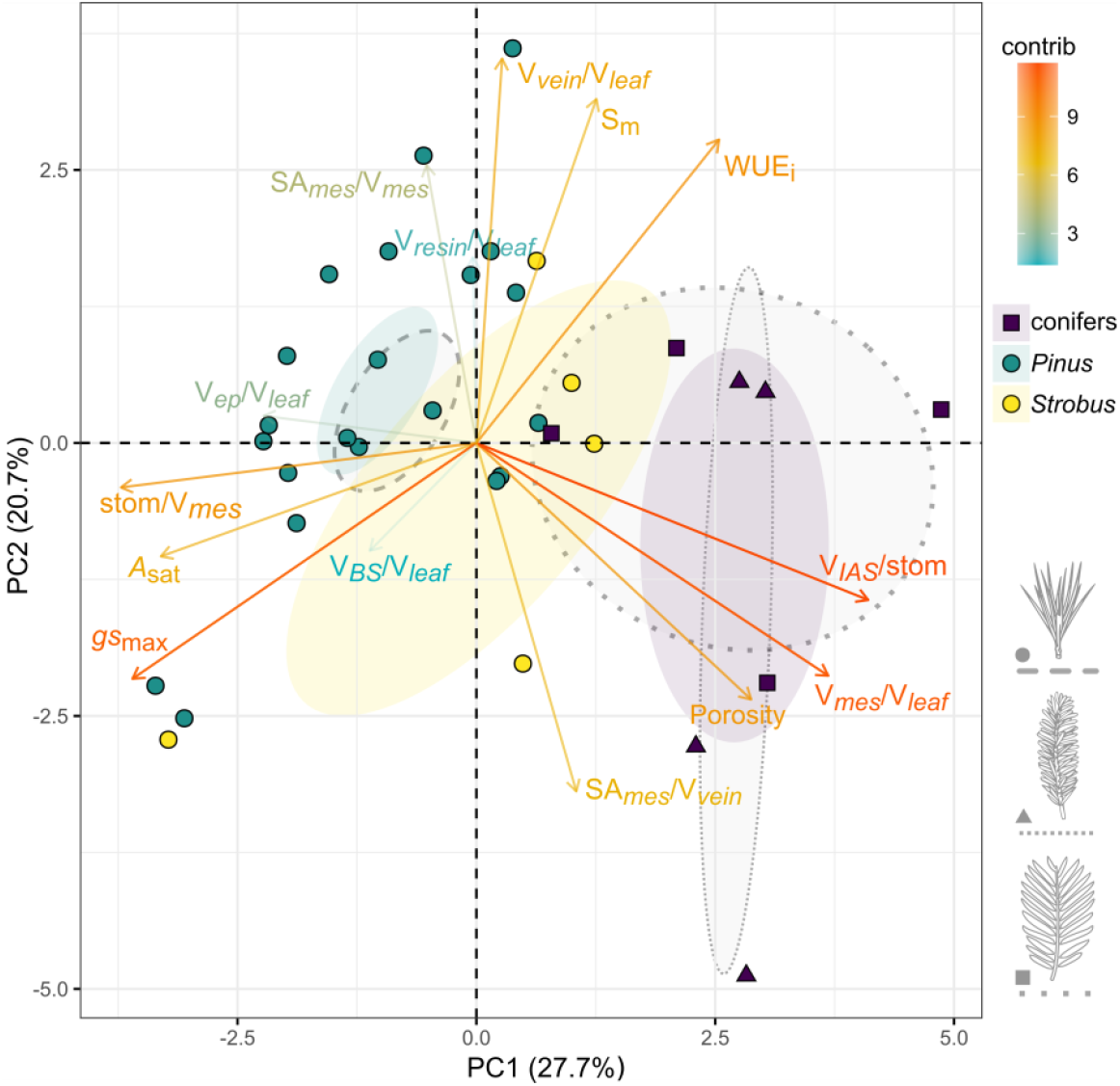
Principal component analysis (PCA) of physiological and volumetric anatomy traits of conifer species. Arrows and trait color gradients indicate the contribution of each variable to the axes. Species from the two *Pinus* subgenera, along with other studied conifer species are indicated in different colors, and 95% confidence ellipses are included. Confidence ellipses for different leaf morphologies are also included. Species bearing flat leaves (squares, dotted line), flattened needle leaves (triangles, narrow dotted line), and needle-like leaves (circles, dashed line) are identified.

**Figure 3.**
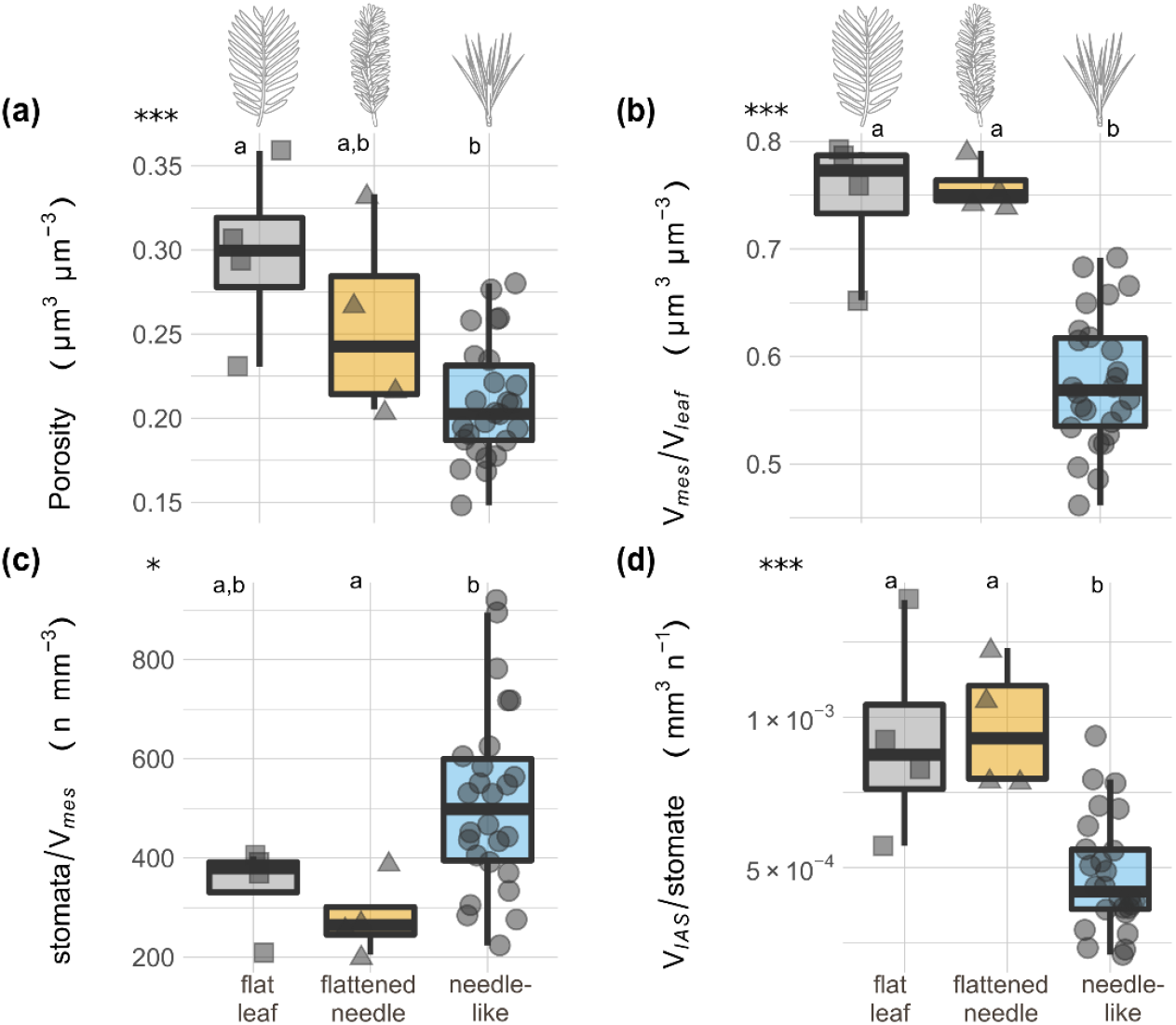
Differences in mesophyll volumetric traits (a,b) and stomatal features (c,d) across flat (squares), flattened needle (triangles) and needle-like (circles) leaves in the conifer species studied. Boxes and bars show the median, quartiles, and extreme values. Gray dots are species data points. *P*-value notations represent results of one-way ANOVAs between groups. * *p* ≤ 0.05; *** *p* ≤ 0.001. Letters indicate significant differences between leaf morphological types. Further information and other comparisons of anatomical traits across leaf morphology types are available in Table S3.

However, flattened needles were differentiated due to some species having significantly higher SA_*mes*_ /V_*vein*_ (Fig. 2; Table S3). Conifer groups were also effectively segregated based on their volumetric anatomy. Non-*Pinus* conifer species were segregated on the first axis (Fig. 2) due to their higher porosity (Fig. 1d), V_*mes*_ /V_*leaf*_, and V_*IAS*_ /stomate values, while species from the *Pinus* subgenus were grouped (Fig. 2) due to higher V_*ep*_ /V_*leaf*_ (Fig. 1b), along with a relatively larger SA_*mes*_ /V_*mes*_. Species from the *Strobus* subgenera were located between the previously described conifer groups (Fig. 2). Structural divergences of leaf morphologies, distinguishing three distinct functional groups, along with the segregation of different conifer clades in the PCA analysis were further supported with one-way ANOVA analyses on 3D leaf anatomical traits and stomatal density (Tables S3,S4). Relative vein volume and SA_*mes*_ /V_*mes*_ were highly conserved (Fig. S3; Tables S3,S4), whereas SA_*mes*_ /V_*vein*_, along with traits related to the ratio of stomatal pore number and mesophyll tissue volumes had significant differences across both leaf morphologies and conifer groups (Figs. 3c,d; S3; Tables S3,S4).

To explore the functional implications of 3D tissue content, we determined their relationships to gas exchange parameters such as *A*_sat_, *g*_smax_, and WUE_i_. WUE_i_ was best predicted by V_*vein*_ /V_*leaf*_ (Fig. 4a), where higher V_*vein*_ /V_*leaf*_ enhances leaf WUE_i_. Further, 2D anatomical estimations of the ratio A_*vein*_ /A_*leaf*_ were comparable to V_*vein*_ /V_*leaf*_, extracted using a 3D approach (Notes S1; Fig. S4). The mesophyll surface area exposed to vein volume (SA_*mes*_ /V_*vein*_) was also an accurate predictor and showed a negative relationship with WUEi (Fig. 4b). A positive relationship of WUE_i_ with S_m_ was also found using generalized least-square models corrected for phylogenetic relatedness (PGLS) analyses (Table S5). *A*_sat_ was the physiological trait showing the least linkage with 3D structural traits. However, we found a negative relationship between *A*_sat_ and mesophyll porosity along with V_*mes*_ /V_*leaf*_ (Fig. S5a), suggesting that conifer species with lower relative mesophyll volumes have greater photosynthetic assimilation rates. Additionally, V_*IAS*_ /stomate was also negatively related to *A*_sat_ (Fig. S5b). The number of stomata per unit mesophyll tissue volume predicted *g*_smax_ and WUE_i_ (Fig. 5). For instance, high stomatal densities relative to mesophyll volume (stomata/V_*mes*_) enhanced *g*_smax_ (Fig. 5a) while decreasing WUE_i_ (Fig. 5b). Evolutionary coordination between leaf volumetric anatomy and physiological traits was supported by PGLS analyses (Table S5). Stomatal density measured on a 2D leaf surface area basis (stomata/A_*leaf*_), included here as a reference of a more standard approach, did not relate to any physiological trait (Table S5). Therefore, the interaction of mesophyll volume and stomatal pore numbers emerged as a key trait to explain leaf physiological performance (Fig. 5). A comparison of the studied conifers with published data for other gymnosperm species, along with angiosperms and ferns, showed that conifers had fewer stomata per unit mesophyll volume than angiosperms *sensu largo* (Fig. S6; Dataset S3). However, this difference was less important when considering evergreen angiosperms alone. Conifers showed a similar stomata/V_*mes*_ ratio as other gymnosperm species and ferns (Fig. S6).

**Figure 4.**
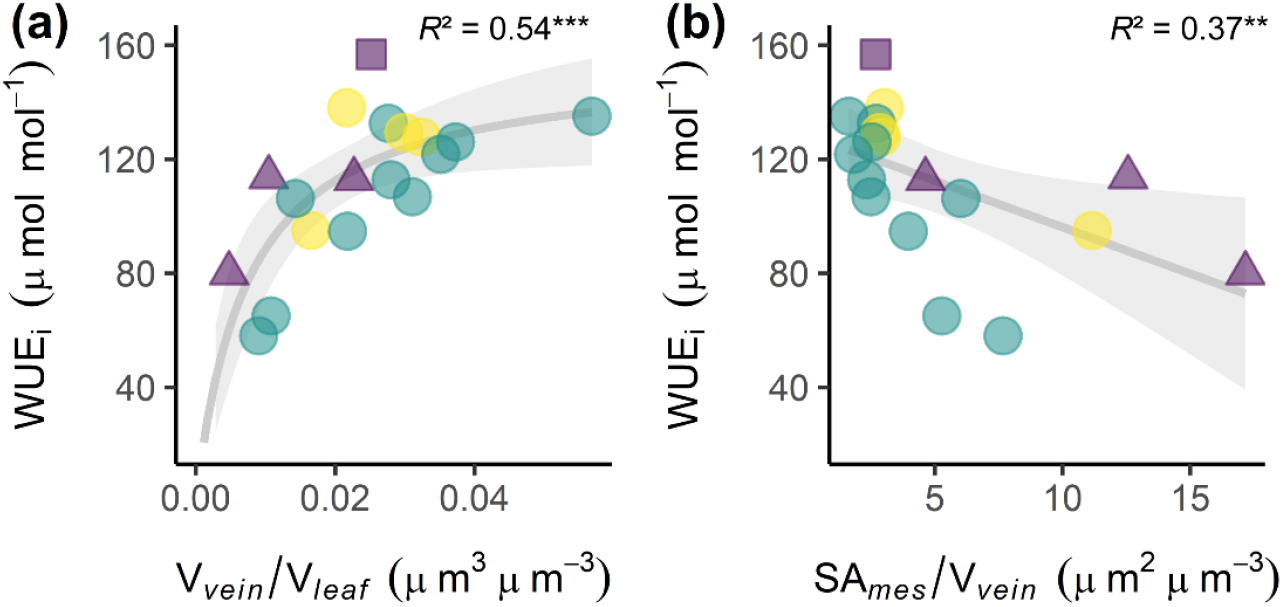
Water use efficiency (WUE_i_) as a function of vein volumetric traits of conifer leaves. Relationships of WUE_i_ with relative vein volume (a) and the surface area of the mesophyll exposed to the IAS per unit of vein volume (b) across 18 conifer species. Solid regression lines and 95% confidence intervals are included. *Pinus* species from the subgenera *Pinus* (green) and *Strobus* (yellow), along with other conifer species (purple) are indicated. Species bearing flat leaves (square), flattened needle leaves (triangles), and needle-like leaves (circles) are also identified. PGLS coefficients of determination are included. ** *p* < 0.01; *** *p* < 0.001. Additional information available in Table S5.

**Figure 5.**
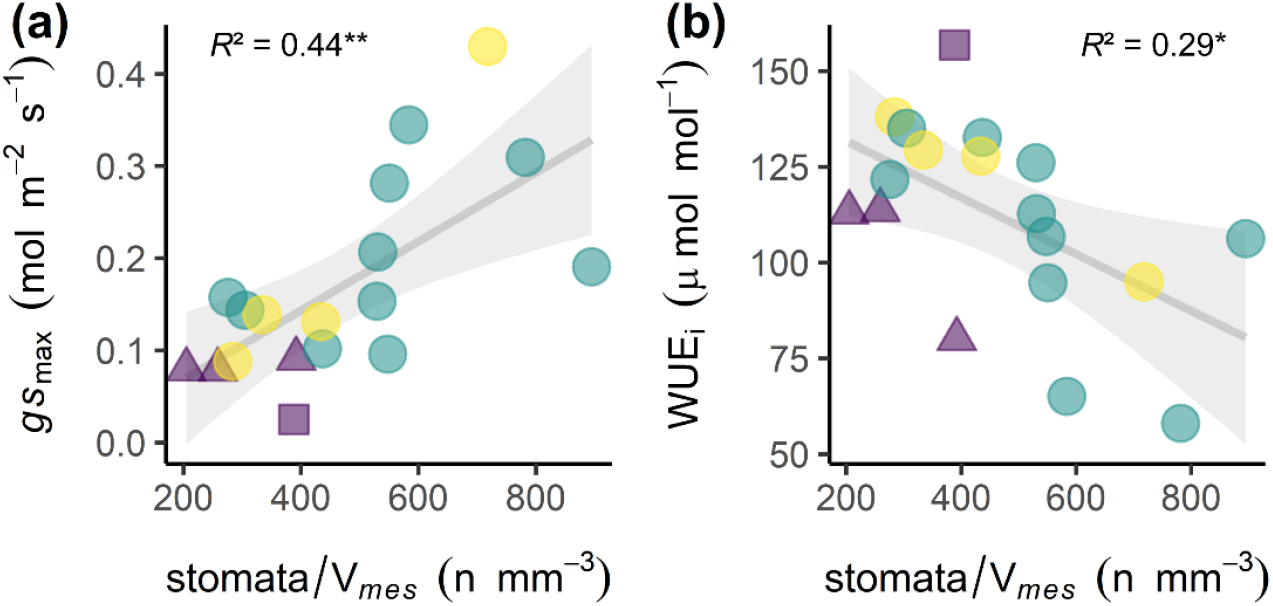
Physiological traits explained by volumetric stomatal density in conifer leaves. Relationships of maximum stomatal conductance (a) and water use efficiency (b) with the number of stomata per unit of mesophyll volume across 18 conifer species. Solid regression lines and 95% confidence intervals are included. *Pinus* species from the subgenera *Pinus* (green) and *Strobus* (yellow), along with other conifer species (purple) are indicated. Species bearing flat leaves (square), flattened needle leaves (triangles), and needle-like leaves (circles) are also identified. PGLS coefficients of determination are included. * *p* < 0.05; ** *p* < 0.01. Additional information available in Table S5.

## Discussion

Conifers, with their long-lasting and anatomically elementary leaves, must rely on leaf construction to enable sufficient carbon assimilation to survive and reproduce while limiting water loss. However, the simple design of coniferous leaves have been considered to have a poor, if not absent, hydraulic connection between the vein xylem and the bulk leaf tissue (Zwieniecki *et al*., 2007). This is unsurprising since conifer leaves are not as fully vascularized as those of angiosperms and present a single cohort of relatively inefficient leaves (Bond, 1989). The likely consequence of this poor hydraulic connection is a larger difference in water potential between the veins and the mesophyll and epidermis in transpiring leaves, which may force stomata to close even when water potential is relatively high in the veins (Zwieniecki *et al*., 2007). Our results show that despite occupying a small fraction (ca. 2%) of the leaf volumetric matrix, vein tissue volume has a great impact on the leaf WUE_i_ (Fig 4a). As the relative vein volume expands, the space between the vascular tissues and the bulk leaf tissues become smaller, potentially reducing the hydraulic resistance for water transport from the vasculature to the bulk leaf. Our study provides an estimation of the full volumetric space occupied by the vein tissue relative to the photosynthetic cells. While we consider volumetric estimates to be more accurate given that they integrate traits over a larger leaf fraction than 2D estimates, we found that standard 2D anatomical estimations of the relative surface covered by veins over a few leaf cross sections could be accurately employed to predict the vein volumetric fraction, and in turn WUE in conifer leaves (Notes S1; Fig. S4). Stomatal and venation densities covary in angiosperms, directly affecting gas exchange and water use capacities (Carins Murphy *et al*., 2014). However, conifers do not provide a dense network of veins within the mesophyll to irrigate the photosynthesizing cells in water. Consequently, conifers might be constrained to increase the relative volume of veins as a single way to provide more water to the leaf.

Previous work has suggested that narrow, needle-like leaves in conifers, which is a common feature in the Pinaceae (Brodribb & Feild, 2008; Brodribb *et al*., 2010), would be a response to alleviate their lack of hydraulic ramification. With the photosynthetic tissue encircling the single vascular cylinder, the distance from the vein to the epidermis largely sets hydraulic conductance outside the xylem, and leaf width becomes a major limiting factor in cylindrical, narrow leaves. Increasing the radial pathlength for water transport should effectively limit leaf hydraulic conductance, in turn limiting photosynthetic rates (Brodribb *et al*., 2010). Our data support this hypothesis, with a negative relationship between mesophyll volume and *A*_sat_ (Fig. S5a). Thus, within our dataset we find evidence for a significant photosynthetic penalty for increasing leaf width and mesophyll volume in flat and flattened needle leaves (Fig. 3b). With a limited ability to maximize photosynthetic capacity through hydraulic ramification, the cylindrical needle-like leaf may offer opportunities for other structural elements that allow improved hydraulic performance. Accessory transfusion tissue is one of them, where specialized cells connect the veins and/or bundle sheath to the mesophyll tissue, providing more water to the photosynthesizing cells and improving hydraulic contact to the epidermis (Brodribb *et al*., 2007). This might lead to increased photosynthetic rates and WUE. Our results support this linkage, with a positive relationship of the combined volumes of bundle sheath and transfusion tissue with photosynthetic assimilation (Table S5). It has been stated that reaching high photosynthetic rates requires high leaf porosity values, which might increase CO_2_ diffusion (Brodribb *et al*., 2020). Yet, we observed a different trend with a negative relationship between mesophyll porosity and *A*_sat_ (Table S5). Moreover, *A*_sat_ was strongly negatively related with V_*mes*_ /V_*leaf*_ in our dataset (Fig. S5a). Previously, a decline of illumination-induced fluorescence as a function of leaf depth was observed in two conifers with needle-like leaves (Johnson *et al*., 2005). Therefore, the decrease of *A*_sat_ in leaves with more voluminous mesophylls might be explained by a limitation of light propagation across the mesophyll. Interestingly, we found conspicuous differences in leaf design between both *Pinus* subgenera and other conifers (Figs. 1,2; Table S4), with narrow needle-leaved *Pinus* possessing less voluminous and porous mesophylls. Such differences in mesophyll construction could explain why *A*_sat_ is greatest in *Pinus* species bearing needle-like leaves.

SA_*mes*_ /V_*vein*_ is another feature that diverged across leaf morphologies and conifer clades (Tables S3,S4). We propose SA_*mes*_ /V_*vein*_ as another anatomical trait involved in regulating the control over the loss of water (Fig. 4b). Minimizing this ratio would mean that less surface is available for evaporation for a given vein water volume, increasing the time before this capacitor is depleted, and thereby lowering the ‘safety margin’ between stomatal closure and xylem cavitation (Zwieniecki *et al*., 2007). Also, in the context of poorly connected hydraulic design, decreasing SA_*mes*_ /V_*vein*_ would minimize the apoplastic surface for water to travel from the vein to the epidermis, hence by proxy decreasing the water path length and increasing connectivity to the epidermis to allow stomata to stay open longer and photosynthesis to continue. In our dataset, decreasing SA_*mes*_ /V_*vein*_ was achieved mainly by increasing the relative volume of veins, i.e. investing more in vascular tissue (Fig. 4). The relative benefit of investing in vein volume to increase the efficiency in water use seems to plateau above ∼3% of leaf volume, and most species produce invests in veins close to that relative volume (Fig. 4a). *Pinus* species, which bear needle-like leaves, have in addition decreased the relative volume of the mesophyll by decreasing airspace volume (Figs. 1; 3a,b; S3a,b) with plicate mesophyll cells (Esau, 1977). Decreasing porosity leads to more cells being in contact with each other, thereby decreasing SA_*mes*_, i.e. the surface of cells exposed to the IAS. Increasing IAS had a positive effect on WUE in six angiosperm species (Mediavilla *et al*., 2001). Although positive, we could not find a significant relationship between mesophyll porosity and WUE_i_ in conifer leaves (Table S5). Beyond considering mesophyll features alone, mesophyll volumetrics in interaction with stomatal pore number emerged here as key traits to explain conifer gas exchange and WUE_i_ (Fig. 5). Using *Arabidopsis* and wheat as model plants, it has been suggested that stomatal differentiation during leaf development might induce mesophyll airspace formation (Lundgren *et al*., 2019). Our study shows that this coordination between V_*IAS*_, along with V_*mes*_, and stomatal number have a significant impact on carbon assimilation and gas exchange on conifers (Fig. 5; Table S5), further supporting this crucial linkage. Considered in a wider context, our observations might provide a novel structural basis to explain the lower photosynthetic rates of ferns and gymnosperms as compared to angiosperms, since we show that they have greater mesophyll volume per stoma, acting as a bottleneck that limits their evaporative capacities (Fig. S6).

Current increases in temperature and atmospheric CO_2_ concentrations might impact the structure and function of conifer forests worldwide, and it has been posed that improved WUE could alleviate the temperature effect (Brodribb *et al*., 2020). Previous studies have shown higher WUE_i_ under increasing CO_2_ in conifer species, having stronger WUE_i_ responses than angiosperms (Dalling *et al*., 2016; Klein *et al*., 2016; Adams *et al*., 2020). Additionally, it has been recently shown that higher plasticity in the vascular tissue of the needles of *Pinus pinaster* enhances their WUE_i_ (Bert *et al*., 2021). Using an experimental approach, needles of *Larix kaempferi* growing under higher CO_2_ showed increased mesophyll surface area per leaf area, coupled with higher photosynthetic rates (Eguchi *et al*., 2004). Moreover, it was shown that elevated CO_2_ increased mesophyll surface and decreased stomatal density in *Pinus sylvestris* needles (Lin *et al*., 2001). Therefore, under such elevated CO_2_ conditions, we might expect to observe a lower stomata/V_*mes*_ ratio, which would enhance WUE_i_ according to our predictions (Fig. 5b). Further, given the recently demonstrated link between enhanced WUEi and vascular tissue plasticity in conifer needles (Bert *et al*., 2021), we expect that coordinated changes in vascular and mesophyll tissue volumetrics, along with shifts in stomatal pore number in conifer leaves, may allow conifer species to cope and adapt to the pressure exerted by increasing VPD in many global biomes (Grossiord *et al*., 2020) by maintaining similar carbon assimilation levels with lower water consumption.

## Acknowledgments

This work was supported by the NSF grants IOS-1626966, IOS-1852976 and IOS-1146746, the Austrian Science Fund (FWF), project M2245, and the Vienna Science and Technology Fund (WWTF), project LS19-013. We thank the Berkeley Arboretum of the University of California Botanical Garden, and the University of Georgia’s Thompson Mills Arboretum for providing plant material. The Advanced Light Source is supported by the Director, Office of Science, Office of Basic Energy Science, of the U.S. Department of Energy under contract no. DE-AC02-05CH11231. TNB acknowledges support from the USDA National Institute of Food and Agriculture (Hatch Award 1016439 and Award 2020-67013-30913) and the NSF (IOS-1951244, IOS-1557906).

## Author Contributions

CB, DMJ, GTR, ST and TNB designed research; CB, DL, DMJ, GTR, JME and ST performed measurements and collected data; ST analyzed the data and wrote the first draft of the manuscript with major contributions from CB and GTR. All authors contributed to manuscript revisions.

## Supporting Information

**The following Supporting Information is available for this article:**

Notes S1. Figures S1 to S6. Tables S1 to S5.

**Other supplementary materials available online:**

Datasets S1 to S3.

### Notes S1. Comparing 2D area fractions to 3D volumetric fractions

Since methods to generate volumetric anatomical data are less available, we were interested in providing a robust alternative using anatomical cross sections, which can be produced using standard light microscopy protocols. Our study shows that V_*vein*_ /V_*leaf*_, the ratio of vein volume over leaf volume, is a powerful explanatory variable and accurately predicts intrinsic water use efficiency (WUE_i_) in conifers (see main text). Given that conifer leaves have a rather homogenous anatomy along their length, we expected that the V_*vein*_ /V_*leaf*_ ratio would be strongly related to the vein surface over leaf surface ratio (A_*vein*_ /A_*leaf*_), which can be measured using standard light microscopy methods on needle cross sections, for example. To test this assumption, we programmatically measured A_*vein*_ /A_*leaf*_ on all slices of each of our scanned and segmented conifer leaf stacks (34 species; total number of slices between ∼200 and 2000) and computed the median value as well as the standard deviation. The relationship between the whole stack median A_*vein*_ /A_*leaf*_ and V_*vein*_ /V_*leaf*_ was very strong (A_*vein*_ /A_leaf_ = 0.0005 + 0.9923 V_*vein*_ /V_*leaf;*_ *R*^2^ = 0.98, *p* < 0.0001). However, we wanted to know if we can reach a similar relationship using the common practice of average anatomical data over a few sections. To do so, we sample one to 50 different slices from the whole stack data, computed the mean A_*vein*_ /A_*leaf*_ from that sample, and repeated that 30 times for each number of slice and for each image stack (Supplementary Figure S4 present data for one to six pooled slices). Using only one slice to estimate A_*vein*_ /A_*leaf*_ give substantially more bias compared to the actual V_*vein*_ /V_*leaf*_ measured, with a larger range of possible values even if the median is close to a 1:1 relationship (A_*vein*_ /A_*leaf*_ = 0.0010 + 0.9576 V_*vein*_ /V_*leaf*_ ; *R*^2^ = 0.86, *p* < 0.0001). Using the mean of three slices produces A_*vein*_ /A_*leaf*_ more comparable to V_*vein*_ /V_*leaf*_, and where 95% of the data falls within one standard deviation of the whole stack A_*vein*_ /A_*leaf*_ data (A_*vein*_ /A_*leaf*_ = 0.0008 + 0.9728 V_*vein*_ /V_*leaf*_ ; *R*^2^ = 0.95, *p* < 0.0001). Averaging over more slices would increase the strength of the relationship. Thus, our analysis shows that area ratios could be used instead of volumetric data. However, as volumetric data provides information over a substantially larger portion of a leaf compared to a few cross sections, it will always provide more accurate data, even in anatomically homogenous leaves such as those of conifer species.

**Figure S1.**
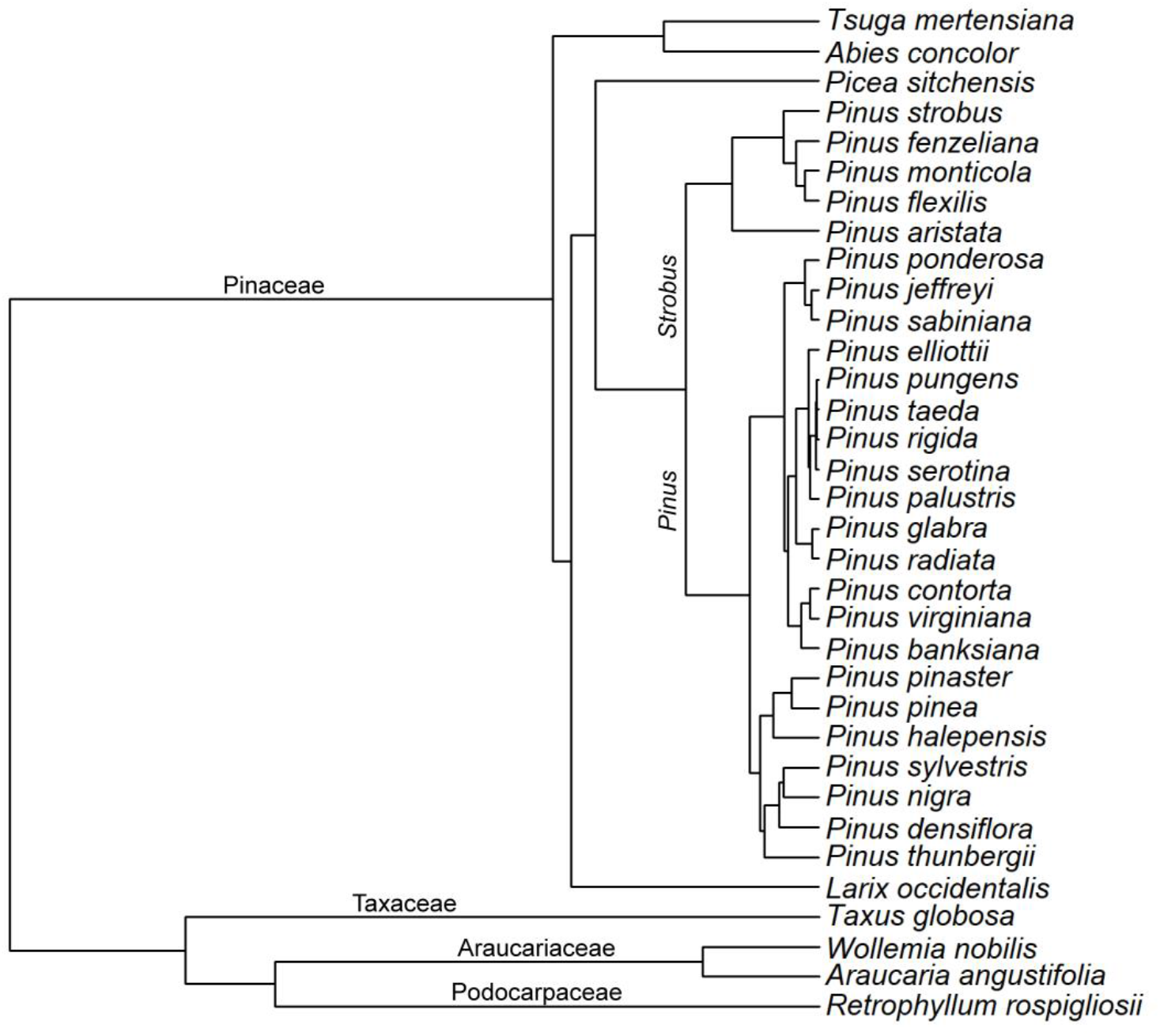
Phylogenetic relationships across the 34 studied species. Labels in branches correspond to different conifer families (horizontal) and *Pinus* subgenera (vertical). The file containing the phylogenetic tree is available as a supplementary material (Dataset S2).

**Figure S2.**
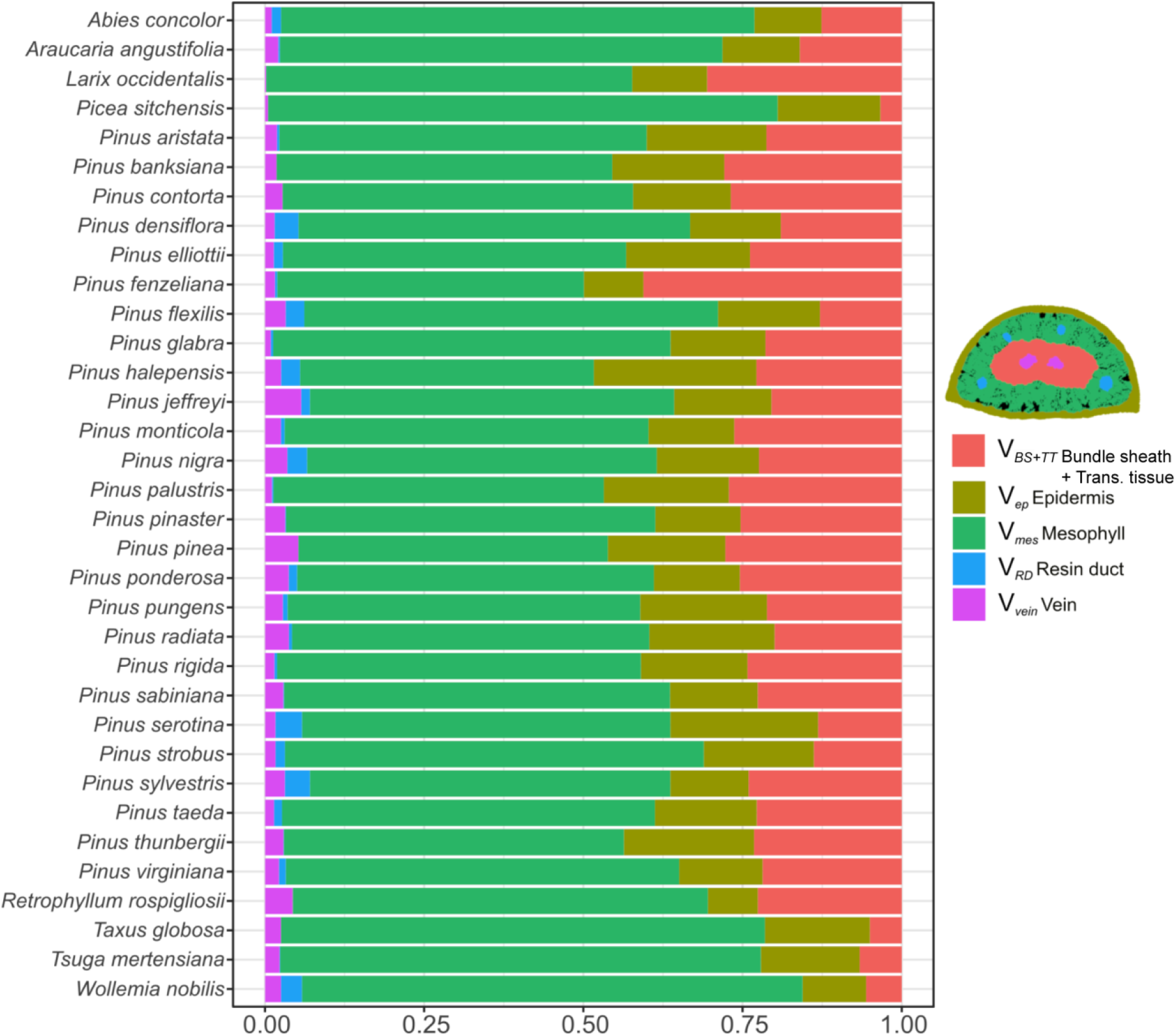
Proportion of tissue relative volumes inside the 3D leaf space for the 34 conifer species studied.

**Figure S3.**
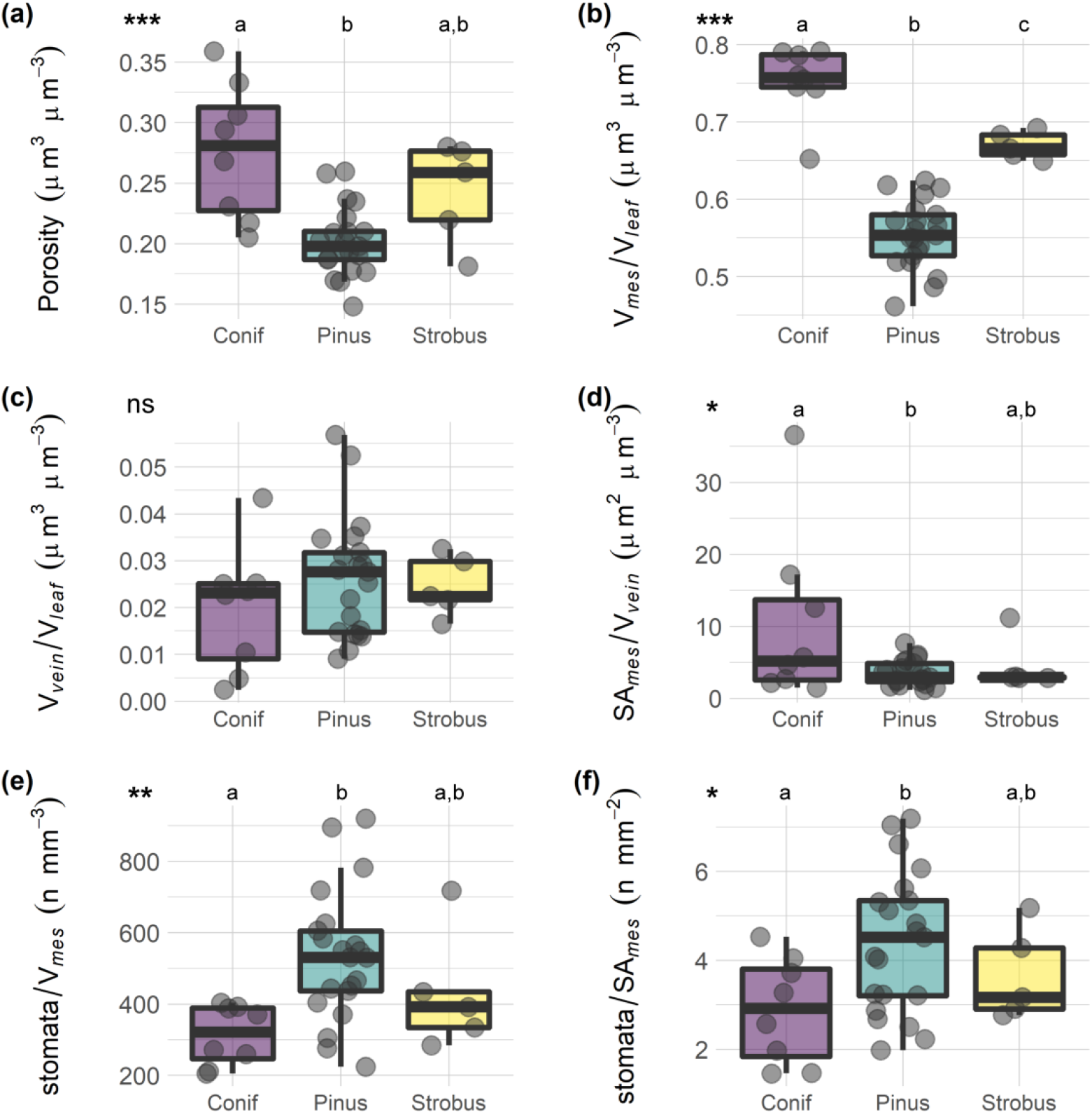
Comparison of leaf 3D structure across conifer groups. Comparison of volumetric anatomy (a-d) and stomatal density (e,f) between the *Pinus* subgenera *Pinus* and *Strobus* and other conifer species studied. Boxes and bars show the median, quartiles, and extreme values. Gray dots are species data points. *P*-values notations represent results of one-way ANOVAs between groups. ns: non-significant; * *p* ≤ 0.05; ** *p* ≤ 0.01; *** *p* ≤ 0.001. Letters indicate significant differences between groups. Further information and other comparisons of traits between groups are available in Supplementary Table 4.

**Figure S4.**
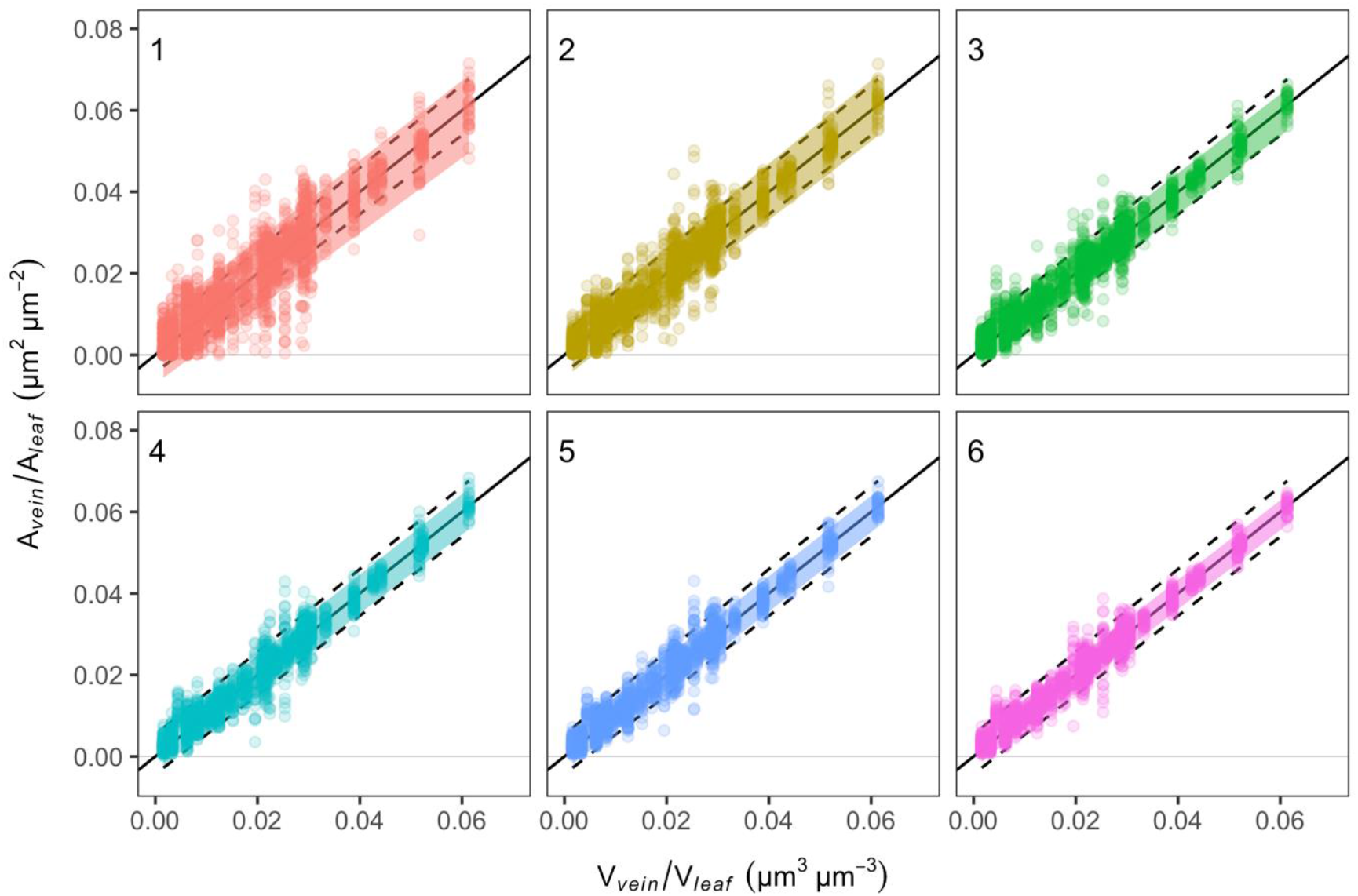
Comparison between 2D area fractions and 3D volumetric fractions in conifer leaf veins. Relationship between vein volume over leaf volume ratio (V_*vein*_ /V_*leaf*_), measured over the whole image stack, and the vein area over leaf area ratio (A_*vein*_ /A_*leaf*_), measured on single image slices of the stacks. In each panel, points represent the mean value of A_*vein*_ /A_*leaf*_ for one to six slices sampled from data collected on each slices of the stack (numbers upper left of the panels represent the number of slices averaged over). Colored ribbons represent the region where 95% of the data lie, and dotted lines represent the A_*vein*_ /A_*leaf*_ median ± standard deviation for the whole stack (measured from ∼200-2000 slices depending on the stack). See the Supplementary Text S1 for additional details.

**Figure S5.**
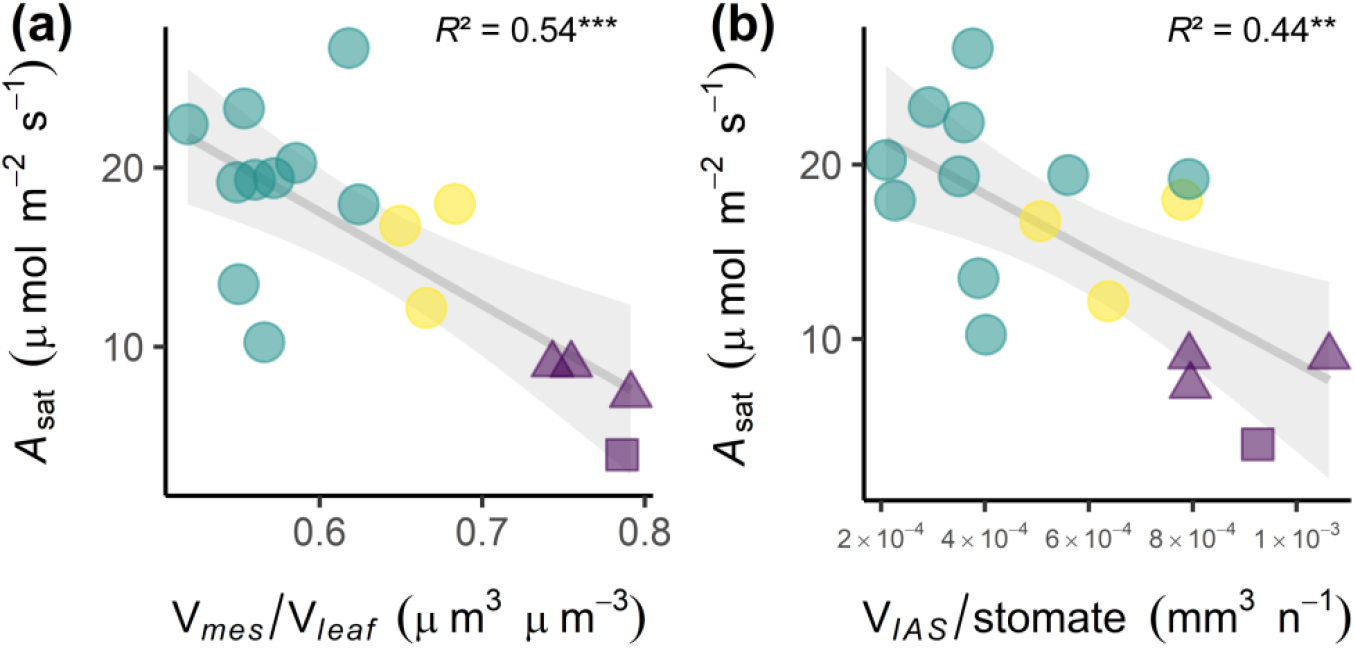
Light-saturated photosynthetic assimilation (*A*_sat_) as a function of volumetric anatomy and stomatal traits in conifers. Relationships of *A*_sat_ with relative mesophyll volume (a) and inter-cellular air space volume (IAS) per stomatal number (b). Solid regression lines and 95% confidence intervals are included. *Pinus* species from the subgenera *Pinus* (green) and *Strobus* (yellow), along with other conifer species (purple) are indicated. Species bearing flat leaves (square), flattened needle leaves (triangles), and needle leaves (circles) are also identified. PGLS coefficients of determination are included. ** *p* < 0.01; *** *p* < 0.001. Additional information available in Supplementary Table 5.

**Figure S6.**
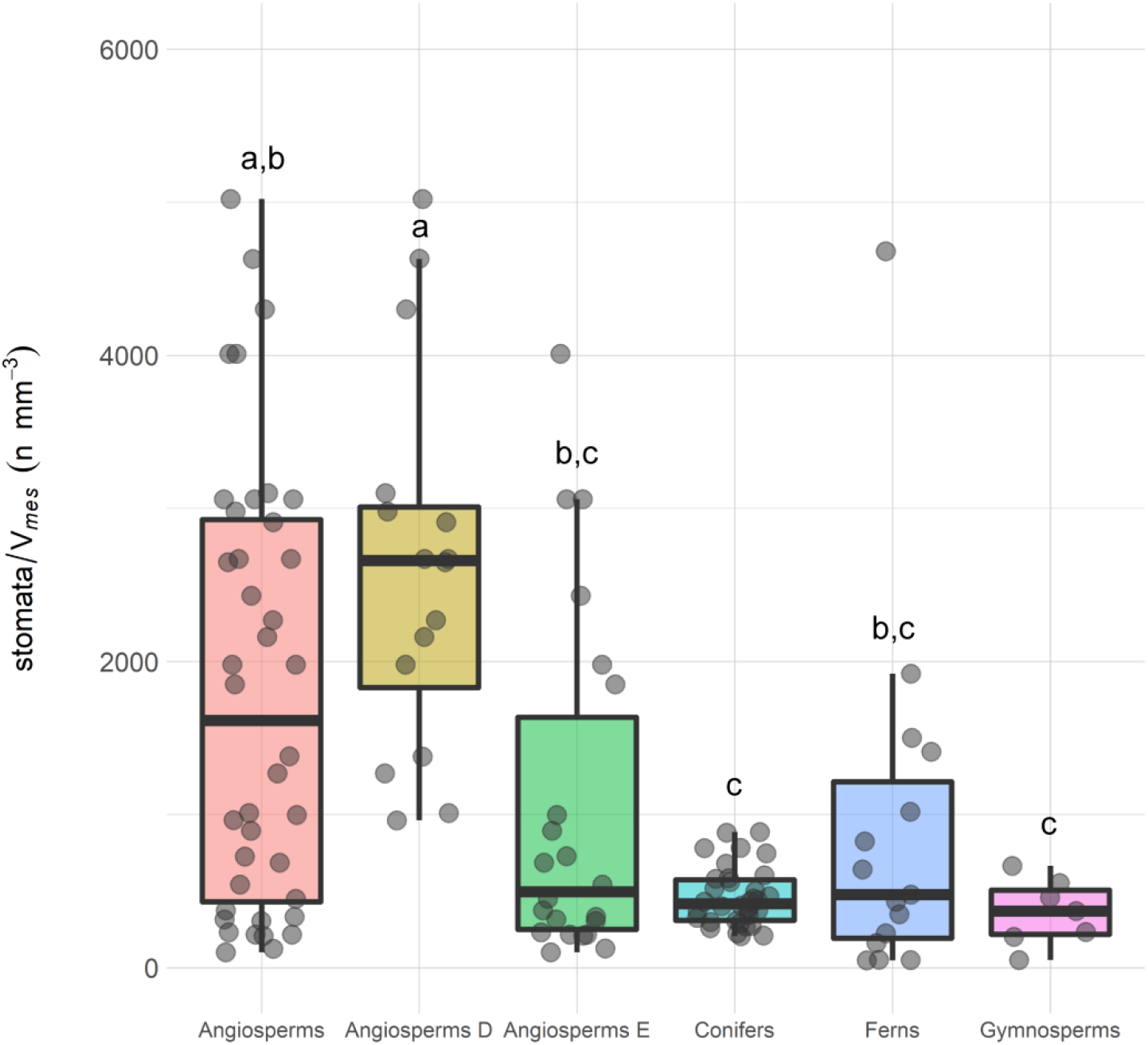
Stomatal number per mesophyll volume across vascular plant groups. Conifer species measured in this study (34 spp) are compared with published data (Théroux-Rancourt *et al*., 2021) for other gymnosperms (7 spp) angiosperms (39 spp) and ferns (16 spp). A distinction between deciduous (D; 17 spp) and evergreen (E; 22 spp) angiosperm species is also included. Significant differences after a one-way analysis of variance followed by a post hoc Tukey’s honest significance differences using 95% confidence intervals are indicated with different letters. The dataset is included as supplementary material (Dataset S3).

**Table S1.**
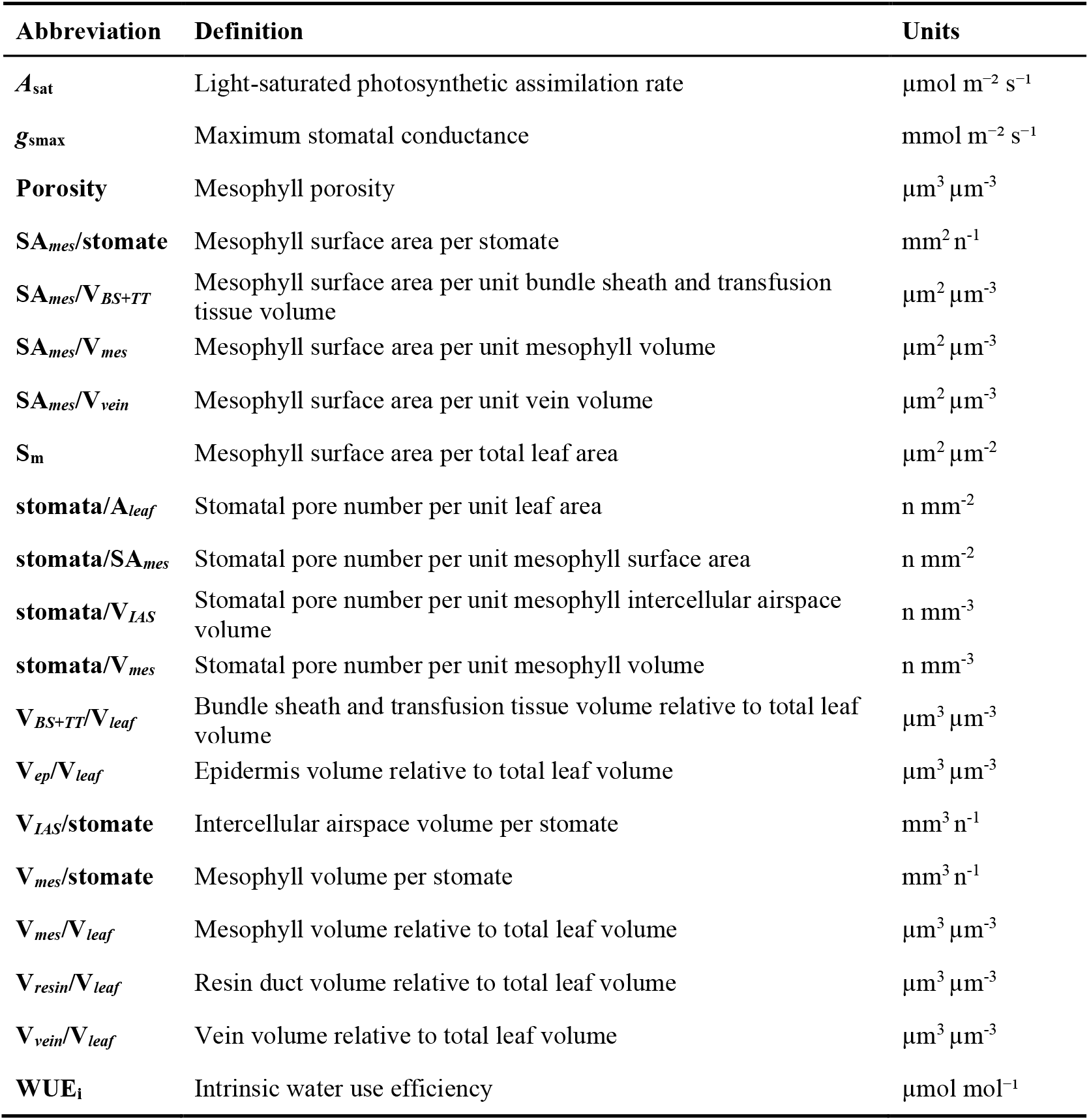
Abbreviations and definitions of anatomical and gas exchange variables measured, with reference to their units.

**Table S2.** Structural coordination within the volumetric space of the conifer leaf. Standardized major axis (SMA) matrix across segmented tissue volumes, relative to total leaf volume, *r*^2^ (above diagonal) and *p*-values (below diagonal) are included. Significant relations at α = 0.05 are indicated.

| | $V_{BS+TT}/V_{leaf}$ | $V_{ep}/V_{leaf}$ | $V_{mes}/V_{leaf}$ | $V_{resin}/V_{leaf}$ | $V_{vein}/V_{leaf}$ |
| --- | --- | --- | --- | --- | --- |
| $V_{BS+TT}/V_{leaf}$ | | 0 | 0.07 | 0.05 | 0 |
| $V_{ep}/V_{leaf}$ | 0.84 | | **0.27** | 0 | 0.01 |
| $V_{mes}/V_{leaf}$ | 0.12 | **<0.01** | | 0.03 | 0.11 |
| $V_{resin}/V_{leaf}$ | 0.19 | 0.93 | 0.33 | | 0 |
| $V_{vein}/V_{leaf}$ | 0.72 | 0.52 | 0.07 | 0.90 | |

**Table S3.** Comparisons of volumetric anatomy and stomatal traits across different leaf morphologies in conifers. Average values for each leaf morphological category are included. Significant differences detected with a one-way ANOVA followed by post hoc Tukey’s honest significance differences tests are indicated with different letters. Significant differences between leaf morphologies at α = 0.05 are also indicated in bold. Graphic representations of some focal comparisons between leaf morphological types are presented in Fig. 3.

| Trait | Flat leaf | Flattened needle | Needle-like | Sum of squares | Mean square | F value (2,31) | Significance |
| --- | --- | --- | --- | --- | --- | --- | --- |
| **Porosity** | 0.30<sup>a</sup> | 0.26<sup>a,b</sup> | 0.21<sup>b</sup> | 0.03081 | 0.015405 | 9.758 | **0.000516** |
| **SA<sub>mes</sub>/stomate** | 0.32<sup>a,b</sup> | 0.51<sup>a</sup> | 0.27<sup>b</sup> | 0.1954 | 0.0977 | 6.708 | **0.0038** |
| **SA<sub>mes</sub>/V<sub>BS+TT</sub>** | 1.15<sup>a</sup> | 1.41<sup>a</sup> | 0.35<sup>b</sup> | 5.493 | 2.7467 | 10.05 | **0.000433** |
| SA<sub>mes</sub>/V<sub>mes</sub> | 0.11<sup>a</sup> | 0.13<sup>a</sup> | 0.12<sup>a</sup> | 0.001002 | 0.0005012 | 0.52 | 0.599 |
| SA<sub>mes</sub>/V<sub>vein</sub> | 3.01<sup>a</sup> | 17.74<sup>b</sup> | 3.69<sup>a</sup> | 707.5 | 353.8 | 16 | **1.69e-5** |
| S<sub>m</sub> | 12.49<sup>a</sup> | 12.75<sup>a</sup> | 10.84<sup>a</sup> | 19.4 | 9.709 | 0.448 | 0.643 |
| **stomata/SA<sub>mes</sub>** | 3.46<sup>a,b</sup> | 2.30<sup>a</sup> | 4.26<sup>b</sup> | 14.23 | 7.117 | 3.448 | **0.0444** |
| **stomata/V<sub>IAS</sub>** | 1190<sup>a</sup> | 1111<sup>a</sup> | 2533<sup>b</sup> | 11710794 | 5855397 | 6.553 | **0.00423** |
| **stomata/V<sub>mes</sub>** | 343<sup>a,b</sup> | 282<sup>a</sup> | 515<sup>b</sup> | 258903 | 129451 | 4.566 | **0.0183** |
| V<sub>BS+TT</sub>/V<sub>leaf</sub> | 0.13<sup>a</sup> | 0.15<sup>a</sup> | 0.24<sup>a</sup> | 0.05873 | 0.029364 | 3.221 | 0.0536 |
| V<sub>ep</sub>/V<sub>leaf</sub> | 0.12<sup>a</sup> | 0.14<sup>a,b</sup> | 0.17<sup>b</sup> | 0.009345 | 0.004673 | 4.654 | **0.0171** |
| V<sub>mes</sub>/V<sub>leaf</sub> | 0.75<sup>a</sup> | 0.76<sup>a</sup> | 0.58<sup>b</sup> | 0.1912 | 0.09560 | 27.56 | **1.32e-7** |
| V<sub>resin</sub>/V<sub>leaf</sub> | 0.009<sup>a</sup> | 0.004<sup>a</sup> | 0.01<sup>a</sup> | 0.000266 | 0.0001331 | 0.807 | 0.456 |
| V<sub>vein</sub>/V<sub>leaf</sub> | 0.03<sup>a</sup> | 0.01<sup>a,b</sup> | 0.02<sup>a</sup> | 0.000973 | 0.0004863 | 3.791 | **0.034** |
| **V<sub>IAS</sub>/stomate** | 0.00092<sup>a</sup> | 0.00097<sup>a</sup> | 0.00047<sup>b</sup> | 1.389e-6 | 6.946e-07 | 15.52 | **2.14e-5** |
| **V<sub>mes</sub>/stomate** | 0.0031<sup>a,b</sup> | 0.0039<sup>a</sup> | 0.0022<sup>b</sup> | 1.096e-5 | 5.481e-6 | 7.392 | **0.00237** |

**Table S4.** Comparisons of volumetric anatomy and physiological features between *Pinus* subgenera *Pinus* and *Strobus*, and other conifer species. Average values for each conifer group are included. Significant differences detected with a one-way ANOVA followed by post hoc Tukey’s honest significance differences tests are indicated with different letters. Significant differences between conifer groups at α = 0.05 are also indicated in bold. Graphic representations of some focal comparisons are presented in Supplementary Fig. 3. *Physiological features measured on a subset of 18 species, different degrees of freedom (2,15) apply to these specific comparisons.

| Trait | *Pinus* | *Strobus* | Other conifers | Sum of squares | Mean square | *F* value (2,31) | Significance |
| --- | --- | --- | --- | --- | --- | --- | --- |
| **Porosity** | 0.20<sup>a</sup> | 0.24<sup>a,b</sup> | 0.28<sup>b</sup> | 0.03438 | 0.017191 | 11.75 | **0.00016** |
| **SA<sub>mes</sub>/stomate** | 0.27<sup>a</sup> | 0.29<sup>a,b</sup> | 0.41<sup>b</sup> | 0.1265 | 0.06323 | 3.766 | **0.034** |
| **SA<sub>mes</sub>/V<sub>BS+TT</sub>** | 0.32<sup>a</sup> | 0.46<sup>a</sup> | 1.28<sup>b</sup> | 5.447 | 9.911 | 2.7237 | **0.00047** |
| SA<sub>mes</sub>/V<sub>mes</sub> | 0.13<sup>a</sup> | 0.12<sup>a</sup> | 0.12<sup>a</sup> | 0.000502 | 0.0002512 | 0.257 | 0.77 |
| SA<sub>mes</sub>/V<sub>vein</sub> | 3.49<sup>a</sup> | 4.55<sup>a,b</sup> | 10.37<sup>b</sup> | 277.9 | 138.93 | 3.862 | **0.032** |
| S<sub>m</sub> | 11.26<sup>a</sup> | 9.15<sup>a</sup> | 12.62<sup>a</sup> | 37.1 | 18.57 | 0.882 | 0.42 |
| **stomata/SA<sub>mes</sub>** | 4.40<sup>a</sup> | 3.66<sup>a,b</sup> | 2.88<sup>b</sup> | 13.77 | 6.88 | 3.312 | **0.049** |
| **stomata/V<sub>IAS</sub>** | 2719<sup>a</sup> | 1754<sup>a,b</sup> | 1151<sup>b</sup> | 15459478 | 7729739 | 10 | **0.00044** |
| **stomata/V<sub>mes</sub>** | 534<sup>a</sup> | 432<sup>a,b</sup> | 312<sup>b</sup> | 293771 | 146886 | 5.394 | **0.0098** |
| V<sub>BS+TT</sub>/V<sub>leaf</sub> | 0.23<sup>a,b</sup> | 0.28<sup>b</sup> | 0.14<sup>b</sup> | 0.06877 | 0.03438 | 3.911 | **0.031** |
| V<sub>ep</sub>/V<sub>leaf</sub> | 0.17<sup>a</sup> | 0.17<sup>a</sup> | 0.13<sup>a,b</sup> | 0.00829 | 0.004147 | 3.995 | **0.029** |
| V<sub>mes</sub>/V<sub>leaf</sub> | 0.55<sup>a</sup> | 0.67<sup>b</sup> | 0.75<sup>c</sup> | 0.24492 | 0.12246 | 70.59 | **2.88e-12** |
| V<sub>resin</sub>/V<sub>leaf</sub> | 0.01<sup>a</sup> | 0.01<sup>a</sup> | 0.006<sup>a</sup> | 0.000218 | 0.0001088 | 0.653 | 0.528 |
| V<sub>vein</sub>/V<sub>leaf</sub> | 0.03<sup>a</sup> | 0.02<sup>a</sup> | 0.02<sup>a</sup> | 0.000245 | 0.0001224 | 0.807 | 0.456 |
| **V<sub>IAS</sub>/stomate** | 0.00044<sup>a</sup> | 0.00060<sup>a</sup> | 0.00095<sup>b</sup> | 1.494e-6 | 7.470e-7 | 18.05 | **6.33e-6** |
| **V<sub>mes</sub>/stomate** | 0.0021<sup>a</sup> | 0.0026<sup>a,b</sup> | 0.0035<sup>b</sup> | 1.062e-5 | 5.312e-6 | 7.061 | **0.0030** |
| A<sub>sat</sub>* | 19.22<sup>a</sup> | 21.91<sup>a</sup> | 7.32<sup>b</sup> | 520.1 | 260.03 | 5.50 | **0.016** |
| g<sub>smax</sub>* | 0.20<sup>a</sup> | 0.19<sup>a</sup> | 0.07<sup>a</sup> | 0.05194 | 0.025971 | 2.713 | 0.099 |
| WUE<sub>i</sub>* | 105.89<sup>a</sup> | 122.51<sup>a</sup> | 115.94<sup>a</sup> | 877 | 438.3 | 0.632 | 0.54 |

**Table S5.** Variation in gas exchange, photosynthesis and water use efficiency as explained by leaf volumetric anatomy and stomatal traits. Phylogenetic generalized least squares (PGLS) statistics are included and significant relations at α = 0.05 are indicated.

| | $A_{\text{sat}}$ | | | | | $g_{\text{smax}}$ | | | | | $\text{WUE}_i$ | | | | |
| --- | --- | --- | --- | --- | --- | --- | --- | --- | --- | --- | --- | --- | --- | --- | --- |
| | Intercept | Slope | $R^2$ | $P$-value | $\lambda$ | Intercept | Slope | $R^2$ | $P$-value | $\lambda$ | Intercept | Slope | $R^2$ | $P$-value | $\lambda$ |
| **Porosity** | **32.35** | **-79.45** | **0.39** | **<0.01** | **0.00** | 0.23 | -0.29 | 0.02 | 0.58 | 0.00 | 93.61 | 85.66 | 0.03 | 0.50 | 0.00 |
| **$\text{SA}_{\text{mes}}/\text{stom}$** | 12.65 | -8.88e-6 | 0.07 | 0.31 | 0.75 | **0.28** | **-3.31e-7** | **0.27** | **<0.05** | **0.00** | **95.96** | **8.68e-5** | **0.26** | **<0.05** | **0.67** |
| **$\text{SA}_{\text{mes}}/\text{V}_{\text{BS+TT}}$** | **15.66** | **-5.92** | **0.28** | **<0.05** | **0.53** | 0.21 | -0.07 | 0.16 | 0.10 | 0.00 | 113.97 | -3.22 | 0.01 | 0.75 | 0.00 |
| $\text{SA}_{\text{mes}}/\text{V}_{\text{mes}}$ | 13.95 | -43.53 | 0.05 | 0.36 | 0.84 | 0.22 | -0.40 | 0.01 | 0.70 | 0.00 | 96.06 | 125.52 | 0.02 | 0.61 | 0.00 |
| $\text{SA}_{\text{mes}}/\text{V}_{\text{vein}}$ | 11.56 | -0.36 | 0.07 | 0.32 | 0.74 | 0.02 | 0.01 | 0.20 | 0.06 | 0.79 | **144.83** | **-3.78** | **0.37** | **<0.01** | **0.52** |
| $\text{S}_m$ | 10.22 | -0.08 | 0.00 | 0.77 | 0.79 | 0.30 | -0.01 | 0.19 | 0.07 | 0.00 | **86.75** | **3.30** | **0.35** | **<0.01** | **0.65** |
| $\text{stom}/\text{A}_{\text{leaf}}$ | 5.56 | 0.11 | 0.11 | 0.19 | 0.70 | 0.06 | 2.57e-3 | 0.13 | 0.14 | 0.00 | 120.80 | -0.21 | 0.01 | 0.63 | 0.00 |
| $\text{stom}/\text{SA}_{\text{mes}}$ | 5.83 | 1.05e+6 | 0.12 | 0.17 | 0.74 | **0.01** | **3.93e+4** | **0.40** | **<0.01** | **0.00** | **163.10** | **-1.01e+7** | **0.43** | **<0.01** | **0.67** |
| $\text{stom}/\text{V}_{\text{IAS}}$ | 7.77 | 1.88e+6 | 0.14 | 0.14 | 0.52 | **0.05** | **5.05e+4** | **0.29** | **<0.05** | **0.00** | **137.21** | **-1.09e+7** | **0.23** | **<0.05** | **0.00** |
| $\text{stom}/\text{V}_{\text{mes}}$ | 7.52 | 5.85e+6 | 0.04 | 0.43 | 0.71 | **-6.36e-3** | **3.74e+5** | **0.44** | **<0.01** | **0.00** | **146.51** | **-7.38e+7** | **0.30** | **<0.05** | **0.00** |
| $\text{V}_{\text{BS+TT}}/\text{V}_{\text{leaf}}$ | **6.13** | **49.65** | **0.38** | **<0.01** | **0.00** | 0.10 | 0.37 | 0.07 | 0.27 | 0.00 | 112.90 | -5.67 | 0.00 | 0.94 | 0.00 |
| $\text{V}_{\text{ep}}/\text{V}_{\text{leaf}}$ | 4.99 | 33.27 | 0.04 | 0.46 | 0.77 | -0.03 | 1.26 | 0.13 | 0.15 | 0.00 | 137.3 | -164.4 | 0.04 | 0.44 | 0.00 |
| $\text{V}_{\text{mes}}/\text{V}_{\text{leaf}}$ | **48.62** | **-51.82** | **0.54** | **<0.001** | **0.00** | 0.49 | -0.51 | 0.17 | 0.08 | 0.00 | 85.25 | 41.84 | 0.02 | 0.57 | 0.00 |
| $\text{V}_{\text{resin}}/\text{V}_{\text{leaf}}$ | 9.97 | -35.45 | 0.00 | 0.74 | 0.78 | 0.20 | -2.04 | 0.05 | 0.35 | 0.00 | 101.3 | 799.1 | 0.14 | 0.12 | 0.00 |
| $\text{V}_{\text{vein}}/\text{V}_{\text{leaf}}$ | 8.68 | 28.85 | 0.00 | 0.77 | 0.79 | 0.18 | -3.34 | 0.17 | 0.09 | 0.58 | **93.19** | **1442.04** | **0.54** | **<0.001** | **0.63** |
| $\text{V}_{\text{IAS}}/\text{stom}$ | **24.74** | **-1.61e-5** | **0.44** | **<0.01** | **0.00** | **0.31** | **-2.58e-7** | **0.38** | **<0.01** | **0.00** | 88.85 | 4.19e-05 | 0.17 | 0.09 | 0.00 |
| $\text{V}_{\text{mes}}/\text{stom}$ | 14.68 | -1.57e-6 | 0.08 | 0.27 | 0.66 | **0.33** | **-6.43e-8** | **0.39** | **<0.01** | **0.00** | 85.35 | 1.04e-05 | 0.18 | 0.08 | 0.00 |

